# *Acacia sieberiana* (Fabaceae) Attenuates Paracetamol and Bile Duct Ligation-Induced Hepatotoxicity via Modulation of Biochemical and Oxidative Stress Biomarkers

**DOI:** 10.1101/2022.06.22.497272

**Authors:** Miriam Watafua, Jane I. Ejiofor, Aminu Musa, Mubarak Hussaini Ahmad

## Abstract

**Background:** The plant *Acacia sieberiana* (Fabaceae) is traditionally used to manage hepatitis. This research work aims to investigate the hepatoprotective effectiveness of root bark extract of *Acacia sieberiana* (ASE) against paracetamol (PCM) and bile duct ligation (BDL)-induced hepatotoxicity. The phytochemical and median lethal dose (LD_50_) investigations were conducted. The rats were pre-treated with the ASE (250, 750, 1,500 mg/kg) once daily via oral route for 7 consecutive days. On the 8^th^ day, liver injury was initiated by PCM administration (2g/kg). Similarly, in the BDL-induced liver injury, the animals were administered ASE (125, 250 and 380 mg/kg) intraperitoneally for 7 consecutive days. After 24 hours, blood samples and hepatic tissues were obtained for biochemical and histopathological investigations.

**Results:** Phytocomponents determination revealed glycosides, triterpenes, glycosides, saponins, tannins, flavonoids and alkaloids. The oral and intraperitoneal LD_50_ values of the ASE were >5,000 and 1,300 mg/kg, respectively. The ASE efficiently (*p*<0.05) decreased the alanine transaminase (ALT) and aspartate transaminase (AST) levels and elevated the albumin and total protein (TP) levels. The direct bilirubin effectively (*p*<0.05) decreased at 750 mg/kg. Besides, the extract efficiently elevated the glutathione peroxidase (GPx), superoxide dismutase (SOD), and catalase (CAT) in relation to the PCM hepatotoxic group. Also, the malondialdehyde (MDA) concentration was reduced by the ASE. Meanwhile, in the BDL– induced liver injury, the ASE remarkably (*p*<0.05) declined the AST, ALP, bilirubin and MDA. Besides, there was effective (*p*<0.05) elevation in SOD, GPx and CAT in the ASE-treated groups. The morphology of liver tissue was preserved at 125 and 250 mg/kg ASE groups from BDL-induced necrosis and vascular congestion.

**Conclusion:** The study shows that the ASE has hepatoprotective actions against liver damage by possible modulation of biochemical and oxidative stress biomarkers

## 1.0 Background

The liver is the largest and one of the most essential organs in the human body, weighing approximately 2–3% of the average body weight (Pingili et al., 2019; Yue et al., 2020). It participates in a lot of physiological activities to keep the normal body system in healthy conditions, such as carbohydrate, lipids and protein metabolism; removal of toxic agents and pathogens; immunological, digestive, nutritional, and storage functions (Duncan et al., 2009; Elmasry, 2021; H. Khan et al., 2019; Protzer et al., 2012). Besides, the liver is the most important location for the biotransformation of exogenous and endogenous chemical compounds (H. Khan et al., 2019). It also participates in key biochemical processes in the body such as growth, energy production, reproduction, maintaining blood glucose level in the fasting stage by gluconeogenesis and glycogenolysis; supply of energy to muscle and brain during starvation; and synthesis of blood clotting factors (A. Khan et al., 2011; Pingili et al., 2019).

As a result of its diverse functions, the liver is frequently vulnerable to both direct and indirect toxic agents (Uzunhisarcikli & Aslanturk, 2019). Hepatic disorders, including cirrhosis, hepatitis, fibrosis and hepatocellular carcinoma, are among the main health care obstacles globally (Elufioye & Habtemariam, 2019). Some factors associated with the pathogenesis of liver disease include lipid peroxidation, reactive oxygen species (ROS), complement factors and pro-inflammatory mediators (chemokines and cytokines) (Elufioye & Habtemariam, 2019). Although the liver has a natural capability to regenerate its lost tissues, certain hepatic injuries or diseases sometimes tend to progress beyond this ability and may result in liver failure or death (Michalopoulos, 2017). Generally, the undiagnosed or unmanaged hepatic injury may progress from acute hepatitis of mere inflammatory reactions to chronic fibrosis, cirrhosis or liver failure (Tag et al., 2015). Besides, it eventually affects the biological functions of body organs (Younossi et al., 2011). Acute or chronic liver disorders lead to high morbidity and cause about 2 million global deaths annually (Asrani et al., 2019). The development and progression of hepatic diseases could be implicated by viruses (hepatitis A, B and C), excessive alcohol intake, malnutrition, and metabolic disorders (Lin et al., 2017; Miltonprabu et al., 2016). Besides, drug-induced hepatic damage serves as the second leading cause of the acute hepatic disorder (Bernal et al., 2010). Some common drugs implicated in liver diseases include paracetamol (PCM), anti-infective agents, anticonvulsants and anti-inflammatory agents (Elufioye & Habtemariam, 2019). Despite the advancement in orthodox medical practice, no effective medications absolutely protect the liver against damage, stimulate its functions, or enhance its hepatic cell regeneration (Madrigal-santillán et al., 2014). Therefore, it is vital to investigate more effective and less toxic alternative therapies to manage liver diseases (Abenavoli et al., 2018; Ma et al., 2020).

Herbal preparations have gained attention in traditional practice against many diseases (Usman et al., 2021). Approximately 75% of the global personalities use herbal preparations for their basic health needs (Muhammad et al., 2021). The biological screening of medicinal plants has motivated the discovery of noble and effective agents against various disorders (Ahmad et al., 2020). About one-third of the medicinal products used in modern medicine were obtained from medicinal plants (Usman et al., 2021). Besides, herbal preparations have gained popularity in traditional practice for their therapeutic uses against hepatic diseases, which stimulates interest in exploring complementary and alternative medicine to develop therapeutically effective compounds against hepatic disorders (Ali et al., 2019; Miltonprabu et al., 2016). Biological investigations have validated the liver protective effect of medicinal plants (Anyasor et al., 2020; Park et al., 2019). Similarly, natural antioxidant compounds have gained attention for utilization against hepatic ailments by virtue of the role played by oxidative stress in the pathogenesis of hepatic disease (Rašković et al., 2014).

The plant *Acacia sieberiana* var Woodii (Fabaceae) is a tree of 3-25 m in height and 0.6-1.8 m in diameter (Ngaffo et al., 2020). The bark is yellowish in colour and rough with gummy exudates. The leaves are sparse and hairy, while the flowers are cream, white or pale yellow (Dawurung et al., 2012). The plant possesses dehiscent shiny brown fruits of approximately 1.3 cm in thickness, 9-21 cm in length and 1.7-3.5 cm in diameter (Dawurung et al., 2012). It is frost and drought resistant and grows in the savannah area with many botanical structures throughout the Sahel and other African nations (Ameh & Eddy, 2014). It is commonly distributed in Ethiopia, Benin, Chad, Gambia, Cameroon, Ghana, Kenya, Liberia, Zimbabwe, South Africa, Mozambique, Senegal, Mali, Mauritania, Namibia, Sierra Leone, Swaziland, Sudan, Nigeria, Portugal, Tanzania, Uganda, Zambia, Togo and India (Ameh & Eddy, 2014). In Nigeria, the plant is available as an economic tree in the Northern regions, including Yobe, Jigawa and Sokoto States (Ameh & Eddy, 2014). It is commonly referred to as umbrella thorn/white thorn/paperback thorn/flat-topped thorn or paperback in English, *Farar kaya* in Hausa, *Aluki* or *Sie* in Yoruba, *Siyi* in Igbo, *Daneji* in Fulfulde languages (Ameh & Eddy, 2014; Salisu et al., 2014).

Previous studies on the phytochemical contents in the *Acacia sieberiana* resulted in isolation of gallic acid, kaempferol, ellagic acid, quercetin, isoferulic acid, kaempferol 3-*α*-_L_-arabinoside and quercetin 3-O-*β*-_D_-glucoside (Abdelhady, 2013). Other secondary metabolites including luteolin-7-O-rutinoside, chrysoeriol-7-O-rutinoside, apigenin-7-O-β-_D_-glucopyranoside, chrysoeriol-7-O-*β*-_D_-glucopyranoside, luteolin, luteolin-3′,4′-dimethoxylether-7-O-*β*-_D_-glucoside and sitosterol-3-O-*β*-_D_-glucoside, were isolated from the leaves of the plant (Ngaffo et al., 2020). The stem and leaves of *Acacia sieberiana* dihydroacacipetalin and acacipetalin (Seigler et al., 1975).

The root decoction of *Acacia sieberiana* is utilized in Nigeria to treat hepatic diseases (Ohemu et al., 2014). The bark and stem of the *Acacia sieberiana* are also used to manage jaundice (Dawurung et al., 2012). Despite the ethnomedicinal applications of the *Acacia sieberiana* in folk medicine to manage liver diseases, no study in the literature scientifically documents its ameliorative actions against liver damage. Hence, the current study evaluates the effect of the methanol root bark extract of *Acacia sieberiana* on PCM- and bile duct ligation-induced liver injuries.

## 2.0 Materials and Methods

### 2.1 Plant collection

The plant *Acacia sieberiana* was obtained from Samaru, Kaduna State, Nigeria. It was identified and validated by Mallam Namadi Sanusi at the herbarium unit of Biological Sciences Department, Ahmadu Bello University (ABU), Zaria, Nigeria (Specimen number= 16136).

### 2.2 Animals

Adult Wistar rats (males and females) ranging from 120g to 200g were sourced from the Animal Facility of the Department of Pharmacology and Therapeutics, ABU, Zaria, Nigeria. They were maintained in well-ventilated animal cages (temperature 22 ± 3 ^0^C, relative humidity of 30-70%) and provided with sufficient animal feed (Vital feed, Jos, Nigeria). sufficient water was supplied to the animals *ad libitum*. The rats acclimatized to the laboratory environment for two weeks before the research work commenced. The study was conducted in accordance with the ABU Ethical Committee on Animal Use and Care Research Policy (ABUCAUC/2016/049) and ARRIVE (Animal Research: Reporting of *In Vivo* Experiments) guidelines. On completing the experiment, the rats were anaesthetized, euthanized by cervical dislocation and buried as per the ABU’s guideline for appropriate disposal of experimental animals remain.

### 2.3 Extraction

The fresh root barks of *Acacia sieberiana* were air-dried in a shaded environment to a uniform weight and size reduced with mortar and pestle. Then 2,500 g of the powdered material was soaked in 10 litres of 70%^v^/_v_ methanol in a conical flask for 72-hours with frequent shaking. The mixture was filtered with Whatman filter paper (No. 1). The filtrate was concentrated on a water bath maintained at 50□C to obtain the extract, packaged and labeled as *Acacia sieberiana* extract (ASE). The mixture of the ASE was prepared freshly for each experiment with distilled water.

Then the extractive value of the extract was obtained as follows:

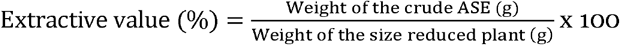

### 2.4 Phytochemical determination

Preliminary phytochemical determination of the ASE was conducted using an appropriate procedure (Sofowora, 1993).

### 2.5 Acute toxicity evaluation

The intraperitoneal (i.p) and oral acute toxicity determination on the ASE were determined in rats in two phases (Lorke, 1983). In the 1^st^ stage, 9 animals were categorized into 3 different groups (n=3), administered the ASE via oral route at 10, 100 and 1,000 mg/kg and then observed for 24-hours for signs of harmful effects or loss of life. In the 2^nd^ stage, 3 rats were categorized into 3 groups (n=1 rat per group) administered with the higher doses of the extract orally (600, 2900 and 5,000 mg/kg) and observed for possible signs of harmful effects and mortality. The same procedure was followed to determine the *i*.*p* acute toxicity. The estimated oral and *i*.*p* median lethal doses (LD_50_) were estimated by taking the geometric mean of the highest non-lethal and the lowest lethal doses as follows:

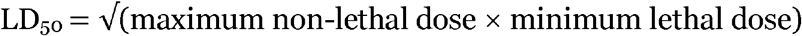

### 2.6 Hepatoprotective effects

#### 2.6.1 Paracetamol-induced hepatotoxicity in rats

The procedure previously described by (Mahmood et al., 2014) was used. The rats were randomly categorised into 6 various groups (n=5) and treated as follows:

**Group 1:** Distilled water (1 ml/kg, *p*.*o*) once daily for 7 days

**Group 2:** Distilled water (1 ml/kg, *p*.*o*) once daily for 7 days

**Group 3:** Silymarin (50 mg/kg, *p*.*o*) once daily for 7 days

**Group 4:** ASE (250 mg/kg, *p*.*o*) once daily for 7 days

**Group 5:** ASE (750 mg/kg, *p*.*o*) once daily for 7 days

**Group 6:** ASE (1,500 mg/kg, *p*.*o*) once daily for 7 days

Following the administration in the above groups (except group 1), liver damage was induced by the PCM (2g/kg) administration (8^th^ day). The rats were anaesthetised with chloroform soaked in cotton, put into an inhalational chamber, and euthanized by cervical dislocation. The blood samples were obtained from the various groups in bottles and centrifuged at 3,000 revolutions per minute (rpm) for 10 minutes. The obtained sera were assayed for biochemical biomarkers such as alanine transaminase (ALT), aspartate transaminase (AST), alkaline phosphatase (ALP), albumin, total protein (TP), total bilirubin (TB) and direct bilirubin (DB). The oxidative stress markers, namely; catalase (CAT), glutathione peroxidase (GPx), superoxide dismutase (SOD), and malondialdehyde (MDA), were also analysed. The livers were excised and placed in 10% formalin for histopathological examination.

#### 2.6.2 Bile duct ligation (BDL)-induced liver injury

The method reported by (Tag et al., 2015) was employed. A total of fifty (50) male animals were anaesthetized with thiopental sodium (40 mg/kg, *i*.*p*). The abdominal furs of the rats were shaved off, and a short incision of about 2 cm was made below the xiphoid process of the abdomen to expose the bile duct region. Double ligation was placed at the common bile duct of forty-four (44) of the rats such that the bile did not flow. The incisions were then sutured, and the animals were kept to recover (fully awake and active). Thirty (30) out of the 44 double ligated rats that were fully awake and active within 24-hours were selected and categorized into 5 groups of 6 rats each, while the 6 non-ligated rats served as a negative control for the study and were treated as follows:

**Group 1 (non-ligated):** Distilled water (1 ml/kg, *p*.*o*) once daily for 7 days

**Group 2:** Distilled water (1 ml/kg, *p*.*o*) once daily for 7 days

**Group 3:** Silymarin (50 mg/kg, *p*.*o*) once daily for 7 days

**Group 4:** ASE (125 mg/kg, *i*.*p*) once daily for 7 days

**Group 5:** ASE (250 mg/kg, *i*.*p*) once daily for 7 days

**Group 6:** ASE (380 mg/kg, *i*.*p*) once daily for 7 days

Then 24-hours post-treatment, the animals were anaesthetised with chloroform and euthanized. The blood samples were obtained from the various groups in bottles and centrifuged at 3,000 rpm. The obtained sera were assayed for biochemical biomarkers such as ALT, AST, ALP, TP, Albumin, TB and DB. The antioxidant markers namely; SOD, CAT, GPx, and MDA, were also assayed. The livers were obtained for histopathological examination.

### 2.7 Data analysis

The values obtained were expressed as mean ± standard error of the mean (SEM) in tables. We used the one way analysis of variance (ANOVA) to analyse the parameters, followed by the Bonferonni post hoc test. The *p*≤0.05 was taken as significant.

## 3.0 Results

### 3.1 Extractive value

A sticky dark-brown solid residue of 115.78g (4.63%^w^/_w_) with a mild sweet smell was obtained from the 2,500g powdered sample of the crude root bark of *A. sieberiana*.

### 3.1 Phytocomponents

The phytocomponents analysis on the ASE indicated cardiac glycosides, saponins, tannins, triterpenes, flavonoids and alkaloids. The steroids and anthraquinones were not present.

### 3.2 Acute toxicity

Acute oral administration of ASE showed no death and behavioural signs of toxicity at 5,000mg/kg, and thus, the oral LD_50_ of the ASE was determined to be ≥5,000 mg/kg. For the i.p acute toxicity study, mortality was not recorded in the first phase of the ASE treatments. However, the rats at all the increased doses (1,600, 2,900 and 5,000 mg/kg) in the second phase died. Hence, the *i*.*p* LD_50_ was about 1,300 mg/kg.

### 3.3 Paracetamol-elicited hepatic injury

#### 3.3.1 Effects of ASE on liver biomarkers of rats in Paracetamol-elicited hepatic injury

The liver of the PCM intoxicated group reflected a remarkable (*p*<0.05) upsurge in ALT, AST and DB and an efficient decrease (*p*<0.05) in TP and albumin when related to the healthy group. The group treated with silymarin and ASE (250 and 750 mg/kg) remarkably reduced the PCM-induced elevated ALT and AST concentrations in relation to the hepatotoxic control group. Besides, the silymarin and ASE at all doses abolished the significant (*p*<0.05) reduction in TP caused by the PCM intoxication. In addition, the extract at all doses efficiently (*p*<0.05) increased the albumin level, while the silymarin pre-treatment showcased a non-significant increase. There was a remarkable (*p*<0.05) decline in the PCM-caused DB elevation in the silymarin and ASE (750 mg/kg) treated groups. The effects of the ASE on the hepatic biomarkers of rats in PCM-elicited liver injury are presented in Table 1.

**Table 1.** Effects of ASE on liver biomarkers of rats in paracetamol-induced liver injury

| Liver biomarkers | Treatment groups (per kg) |  |  |  |  |  |
| --- | --- | --- | --- | --- | --- | --- |
|  | D/W<br>(1 ml) | D/W (1 ml)<br>+ PCM (2 g) | Sily (50mg)<br>+ PCM (2g) | ASE (250mg)<br>+ PCM (2g) | ASE (750mg)<br>+ PCM (2g) | ASE (1,500mg)<br>+ PCM (2g) |
| ALT (IU/L) | 11.33±0.88 | 30.33±2.03<sup>a</sup> | 17.00±1.15<sup>a,b</sup> | 16.33±1.45<sup>a,b</sup> | 17.00±2.65<sup>a,b</sup> | 26.33±3.76<sup>a</sup> |
| AST (IU/L) | 38.67±3.48 | 45.33±0.40<sup>a</sup> | 41.00±3.79<sup>b</sup> | 37.33±1.45<sup>b</sup> | 38.33±4.63<sup>b</sup> | 41.35±4.61 |
| ALP (IU/L) | 12.49±1.18 | 13.58±0.53 | 14.55±1.05 | 14.05±1.16 | 11.70±1.03<sup>a</sup> | 12.64±1.05 |
| Total Protein<br>(g/dL) | 9.37±0.53 | 8.05±0.09<sup>a</sup> | 9.14±0.19<sup>b</sup> | 8.86±0.17<sup>b</sup> | 10.16±0.91<sup>b</sup> | 10.14±0.95<sup>b</sup> |
| Albumin (g/dL) | 3.85±0.81 | 2.64±0.27<sup>a</sup> | 3.03±0.56 | 3.25±0.18<sup>b</sup> | 3.46±0.19<sup>b</sup> | 3.00±0.12<sup>b</sup> |
| Direct bilirubin<br>(mmol/L) | 3.03±0.44 | 4.24±0.68<sup>a</sup> | 3.23±0.21<sup>b</sup> | 3.56±0.43 | 3.18±0.26<sup>b</sup> | 4.07±1.19 |
| Total bilirubin<br>(mmol/L) | 5.30±0.35 | 5.78±0.41 | 5.33±0.43 | 5.45±0.30 | 5.16±0.97 | 5.69±0.46 |

n = 5; The parameters are expressed as mean ± SEM; One Way ANOVA followed by Bonferroni post hoc test; <sup>a</sup> $p < 0.05$ significant compared to the D/W; <sup>b</sup> $p < 0.05$ significant compared to the D/W+PCM group. Sily = Silymarin; D/W= Distilled water; PCM=PCM; ALT=alanine aminotransferase; AST=aspartate aminotransferase; ALP=alkaline phosphatase; ASE = *Acacia sieberiana* extract

#### 3.3.2 Effects of ASE on oxidative stress biomarkers in paracetamol-elicited liver injury

This result showed that the PCM-elicited hepatic damage significantly reduced (*p*<0.05) the SOD level in relation to the healthy rats. However, the groups treated with the silymarin and ASE at all doses efficiently (*p*<0.05) and dose-dependently elevated SOD plasma concentration related to the PCM-hepatotoxic group. The concentration of the CAT was not affected by the PCM intoxication. However, the CAT level significantly increased (*p*<0.05) in the categories that received the ASE at 250 and 750 mg/kg compared to the distilled water healthy and hepatotoxic groups. The PCM intoxicated group elicited significant (*p*<0.05) elevation in the MDA concentration, which was reversed effectively (*p*<0.05) by the silymarin and ASE at all doses. The PCM-induced injury slightly and insignificantly reduced the GPx concentration. However, its serum concentration was increased in all the ASE pre-treated groups in relation to the distilled water healthy group. There was an effective (*p*<0.05) increase in the GPx in the silymarin pre-treated group in relation to the PCM-intoxicated category. The actions of the ASE on the oxidative stress biomarkers of rats in PCM-induced liver injury are presented in Table 2.

**Table 2.** Effects of the ASE on oxidative stress biomarkers of rats in paracetamol-induced liver injury

| Treatment groups<br>(per kg) | Oxidative stress biomarkers |  |  |  |
| --- | --- | --- | --- | --- |
|  | SOD (ng/mL) | CAT (ng/mL) | MDA (nmol/mL) | GPx (ng/mL) |
| D/W (1 ml) | 9.47 ±1.64 | 40.94 ±1.45 | 13.71 ±0.65 | 92.39 ±4.42 |
| D/W (1 ml) + PCM | 6.58 ±0.74<sup>a</sup> | 41.31 ±2.39 | 27.00 ±3.60<sup>a</sup> | 89.62 ±5.80 |
| Sily (50 mg) + PCM | 11.45 ±1.61<sup>b</sup> | 48.90 ±6.64<sup>a</sup> | 14.17 ±1.19<sup>b</sup> | 99.92 ±1.61<sup>a,b</sup> |
| ASE (250) + PCM | 10.71 ±0.81<sup>b</sup> | 45.58 ±1.26<sup>a,b</sup> | 12.28 ±0.50<sup>b</sup> | 99.88 ±9.81 |
| ASE (750) + PCM | 12.55 ±0.81<sup>a,b</sup> | 53.91 ±3.17<sup>a,b</sup> | 11.93 ±0.96<sup>b</sup> | 97.06 ±4.62 |
| ASE (1500) + PCM | 13.22 ±0.97<sup>a,b</sup> | 40.82 ±3.07 | 12.11 ±0.65<sup>b</sup> | 94.59 ±9.77 |

n = 5; parameters are expressed as mean ± SEM; One Way ANOVA followed by Bonferroni post hoc test; <sup>a</sup> $p < 0.05$ significant compared to the D/W; <sup>b</sup> $p < 0.05$ significant compared to the D/W+PCM. Sily = silymarin; D/W= Distilled water; PCM=PCM; SOD=Superoxide dismutase; CAT= Catalase; MDA=Malondialdehyde; GPx=glutathione peroxidase; ASE = *Acacia* *sieberiana* extract

#### 3.3.3 Effects of ASE on the liver histology in the paracetamol-elicited liver injury

The histopathological results of the rats’ livers after PCM-induced liver injuries showed that the PCM produced intense hepatocellular necrosis with sinusoid and vascular congestion. At the same time, the group pre-treated with the standard drug silymarin exhibited slight focal necrosis, lymphocyte hyperplasia and vascular congestion. There was slight kupffer cell hyperplasia and slight sinusoidal congestion in the group that received ASE at 250mg/kg, while the group pre-treated with the ASE at 750mg/kg showed vascular congestion and slight vacuolation and necrosis. However, the group pre-treated with 1,500mg/kg of ASE showed intense hepatocellular necrosis and lymphocyte hyperplasia. The actions of the ASE on the hepatic histology of rats in PCM-produced liver injury are shown in Figure 1.

**Figure 1:**
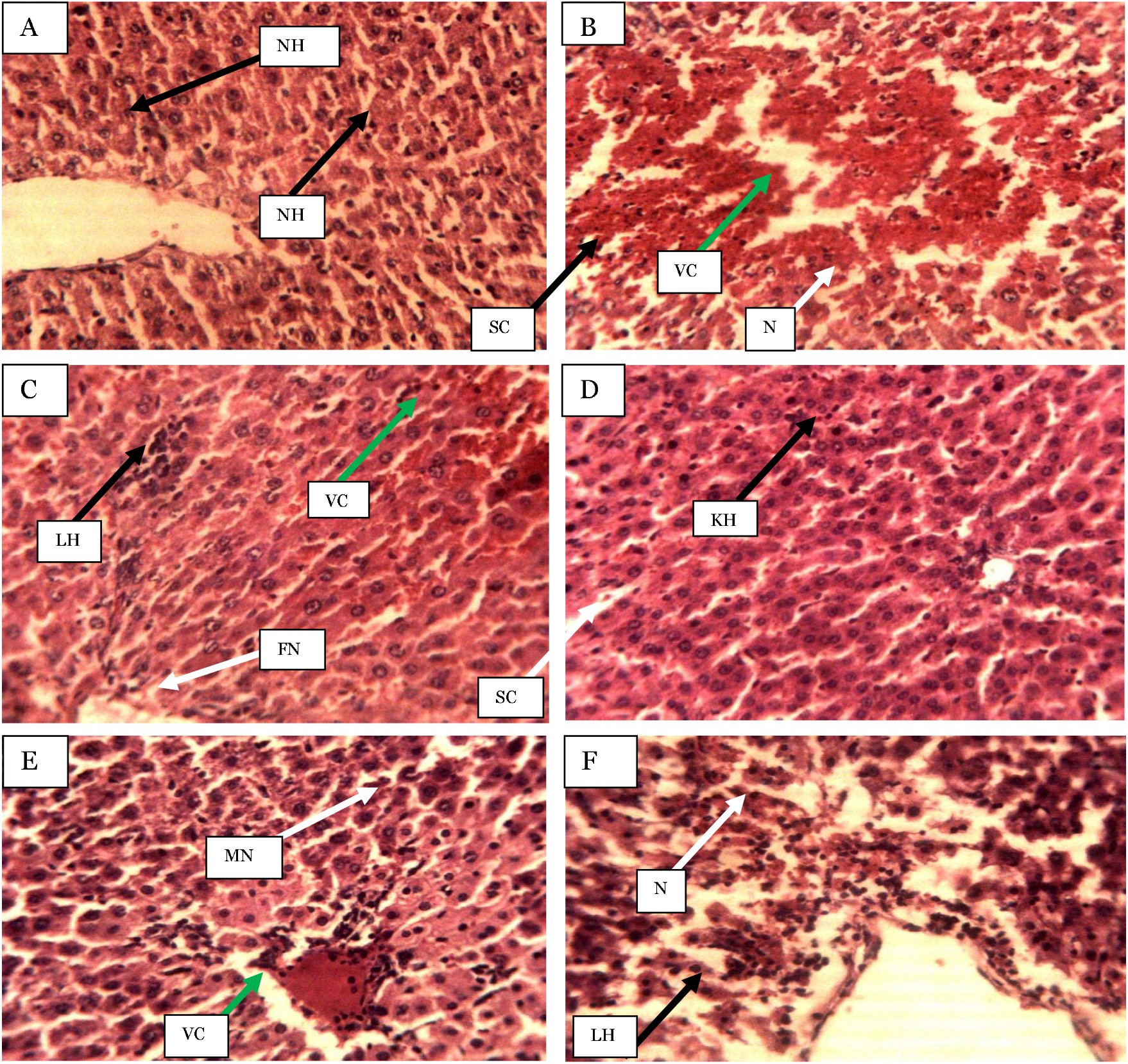
Photomicrographs of hepatic sections of rats after 7-days of ASE pre-treatment in the paracetamol-induced liver injury (Haematoxylin and eosin-stained at ×250 magnification) A) Distilled water (1 ml/kg), B) Distilled water (1 ml/kg) + PCM), C) Sily (50 mg/kg) + PCM), D) ASE (250 mg/kg) + PCM), E) ASE (750 mg/kg) + PCM), F) ASE (1,500 mg/kg) + PCM), NH= Normal hepatocytes, N= Necrosis, SC= Sinusoid congestion, VC= Vascular congestion, FN= Focal necrosis, LH= Lymphocytes hyperplasia, KH= Kupffer cell hyperplasia, SC= slight sinusoidal congestion, MN= Moderate necrosis, ASE = *Acacia sieberiana* extract

### 3.4 Bile duct ligation-elicited hepatic injury

#### 3.4.1 Effects of ASE on hepatic biomarkers of rats in bile duct ligation-induced liver injury

All the liver biomarkers in the BDL-induced hepatic injured group were elevated, which were significant (*p*<0.05) for ALP, AST, and bilirubins in relation to the non-ligated control group. However, the BDL produced a non-significant reduction in the albumin level related to the non-ligated control group. The ASE non-significantly declined the ALT and total protein levels at all doses compared to the BDL-treated control group. The ASE also produced a remarkable (*p*<0.05) reduction in the AST (125 and 250 mg/kg), ALP (250 and 380 mg/kg), direct and total bilirubin levels at all doses. However, the ASE at higher doses (250 and 380 mg/kg) produced a non-significant increase in the plasma albumin concentration. The standard agent silymarin increased the ALT, total protein, and albumin related to the BDL-ligated control category. However, the ALP and AST declined insignificantly in the category pre-treated with the silymarin in relation to the BDL-ligated control group. Also, the direct and total bilirubin reduced (*p*<0.05) in the silymarin administered group. The effects of the ASE on the hepatic biomarkers of rats in BDL-induced liver injury are showcased in Table 3.

**Table 3.** Effects of ASE on liver biomarkers of rats in bile duct ligation-induced liver injury

| Liver biomarkers | Treatment groups (per kg) |  |  |  |  |  |
| --- | --- | --- | --- | --- | --- | --- |
|  | Non-ligated<br>+ D/W (1ml) | BDL +<br>D/W (1ml) | BDL+ Sily<br>(50 mg) | BDL+ASE<br>(125 mg) | BDL+ASE<br>(250 mg) | BDL+ASE<br>(380 mg) |
| ALT (IU/L) | 7.67±0.88 | 8.33±1.45 | 9.02±1.00 | 6.67±0.33<sup>c</sup> | 7.00 ±1.0 2 | 7.67±1.45 |
| AST (IU/L) | 9.00 ±1.15 | 24.67±4.17<sup>a</sup> | 20.00±3.58<sup>a</sup> | 12.05±2.08<sup>b,c</sup> | 10.50±0.50<sup>b,c</sup> | 19.67±1.76<sup>a</sup> |
| ALP (IU/L) | 12.87± 2.04 | 16.23±0.84<sup>a</sup> | 13.32±2.18 | 15.93±0.38<sup>a</sup> | 12.57±0.26<sup>b</sup> | 13.93±1.21<sup>b</sup> |
| Total Protein (g/dL) | 5.98±0.09 | 6.35±0.90 | 6.78±1.16 | 5.39±0.60 | 5.59±0.50 | 5.62±0.64 |
| Albumin (g/dL) | 1.88±0.37 | 1.43±0.38 | 2.20±0.16<sup>b</sup> | 1.42±0.31 | 1.64±0.30 | 1.91±0.13 |
| Direct bilirubin (mmol/L) | 0.33±0.13 | 2.83±0.08<sup>a</sup> | 0.39±0.08<sup>b</sup> | 0.27±0.04<sup>b</sup> | 0.23±0.10<sup>b</sup> | 0.27±0.04<sup>b</sup> |
| Total bilirubin (mmol/L) | 0.56 ±0.17 | 2.86±0.08<sup>a</sup> | 1.18±0.23<sup>a,b</sup> | 0.82±0.27<sup>b</sup> | 0.58±0.10<sup>b</sup> | 1.53±0.10<sup>a,b</sup> |

n = 6; The parameters expressed as mean ± SEM; One Way ANOVA followed by Bonferroni post hoc test; <sup>a</sup> $p < 0.05$ significant compared to the Non-ligated + D/W; <sup>b</sup> $p < 0.05$ significant compared to the BDL + D/W; <sup>c</sup> $p < 0.05$ significant compared to the BDL + Sily; Sily = silymarin; D/W=Distilled water; BDL=Bile duct ligation; ALT=alanine aminotransferase; AST=aspartate aminotransferase; ALP=alkaline phosphatase; ASE = *Acacia sieberiana* extract

#### 3.4.2 Effects of ASE on oxidative stress biomarkers in bile duct ligation-induced liver injury

The SOD and CAT concentration effectively (*p*<0.05) reduced in the BDL-induced liver injured group related to the non-ligated control class. The MDA level was increased significantly (*p*<0.05) in the BDL-induced liver injured group. In addition, the GPx level was slightly elevated in the BDL-ligated group. The standard drug (silymarin) and ASE (125 and 250 mg/kg) showed a remarkable (*p*<0.05) upsurge in the serum SOD concentration in comparison to the BDL-treated control group. Similarly, the silymarin and extract elicited dose-dependent and significant elevation in the CAT level. Also, an efficient (*p*<0.05) increase in the GPx was observed in all the extract-treated groups. The standard agent (silymarin) and ASE (125 mg/kg) showed a remarkable (*p*<0.05) reduction in the MDA serum concentration related to the BDL-treated control group. The effects of the ASE on the oxidative stress biomarkers of rats in BDL-induced liver injury are presented in Table 4.

**Table 4.** Effects of ASE on oxidative stress biomarkers in bile duct ligation-induced liver injury

| Treatment groups<br>(per kg) | Oxidative stress biomarkers |  |  |  |
| --- | --- | --- | --- | --- |
|  | SOD(ng/mL) | CAT (ng/mL) | MDA (nmol/mL) | GPx (ng/mL) |
| Non-ligated +D/W (1 ml) | 5.47 ±0.65 | 40.94 ±1.45 | 36.71 ±1.54 | 95.06 ±4.05 |
| BDL + D/W (1 ml) | 1.27 ±0.18<sup>a</sup> | 32.48 ±8.27<sup>a</sup> | 68.61 ±6.09<sup>a</sup> | 99.61 ±1.02 |
| BDL + Sily (50 mg) | 2.01 ±0.13<sup>a,b</sup> | 61.38 ±5.30<sup>a,b</sup> | 47.30 ±9.44<sup>b</sup> | 105.28 ±7.40 |
| BDL + ASE (125 mg) | 2.35 ±0.40<sup>a,b</sup> | 45.06 ±3.24<sup>b,c</sup> | 52.45 ±3.91<sup>a,b</sup> | 112.62±8.31<sup>a,b</sup> |
| BDL + ASE (250 mg) | 2.61 ±0.31<sup>a,b</sup> | 49.11±4.95<sup>a,b,c</sup> | 67.45 ±7.45<sup>a</sup> | 116.50±7.50<sup>a,b</sup> |
| BDL + ASE (380 mg) | 1.65 ±0.37<sup>a</sup> | 53.70 ±3.75<sup>a,b</sup> | 68.23 ±6.23<sup>a</sup> | 113.54±7.54<sup>a,b</sup> |

n = 6; The values are expressed as mean ± SEM; One Way ANOVA followed by Bonferroni post hoc test; <sup>a</sup> $p < 0.05$ compared to the Non-ligated + D/W; <sup>b</sup> $p < 0.05$ compared to the BDL + D/W, <sup>c</sup> $p < 0.05$ compared to the BDL + Sily group. Sily =Silymarin; D/W= Distilled water; BDL=Bile duct ligation; CAT=Catalase; SOD=Superoxide dismutase; GPx=Glutathione peroxidase; MDA=Malondialdehyde; ASE = *Acacia sieberiana* extract

#### 3.4.3 Effects of ASE on the liver histology in the bile duct ligation-induced liver injury

The hepatocytes of the operated, non-ligated control group were intact with normal structures. However, the BDL-injured group showed moderate hepatic necrosis and vascular congestion, which was slightly reversed in the group that received the ASE at 125mg/kg. The group treated with the extract at 250mg/kg showed lymphocyte hyperplasia. The ASE at the highest dose (380 mg/kg) revealed Kupffer cell hyperplasia and vascular congestion. The effects of the ASE on the liver histology are shown in Figure 2.

**Figure 1:**
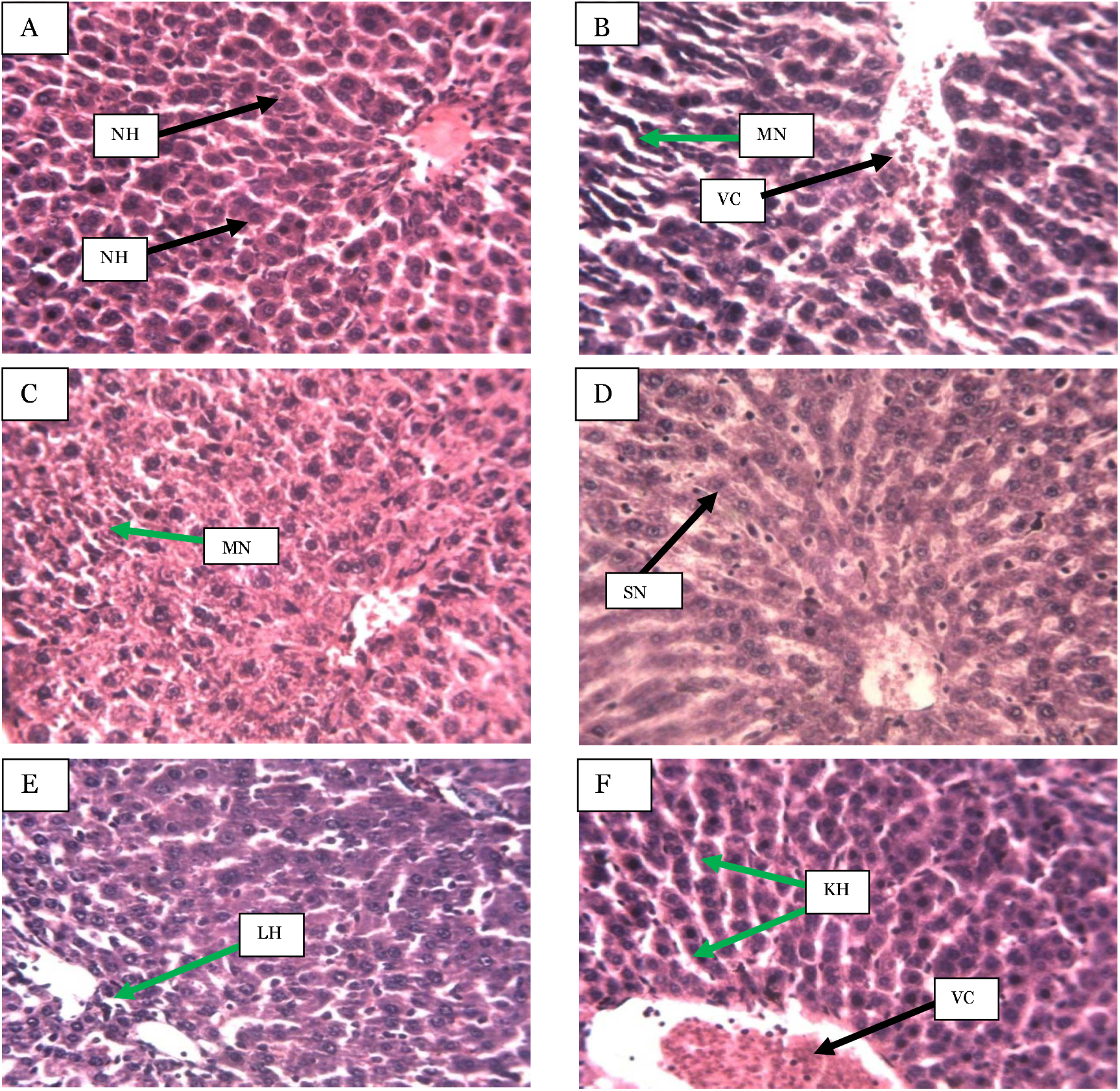
Photomicrographs of hepatic sections of rats after 7-days of ASE treatments in the bile duct ligation-induced liver injury (Haematoxylin and eosin-stained at ×250 magnification) A) Distilled water (1 ml/kg), B) BDL + Distilled water (1 ml/kg), C) BDL + Sily (50 mg/kg), D) BDL + ASE (125 mg/kg), E) BDL + ASE (250 mg/kg), F) BDL + ASE (380 mg/kg), NH= Normal hepatocytes, VC= Vascular congestion, MN= Moderate necrosis, SN= Slight necrosis, LH= Lymphocytes hyperplasia, KH= Kupffer cell hyperplasia, ASE = *Acacia sieberiana* extract

## 4.0 Discussion

The liver is often exposed to diverse endogenous and exogenous toxic xenobiotics, drugs, metabolic waste products, and viral or bacterial agents which traverse it for detoxification or excretion (Zhou et al., 2019). This predisposes the liver to the harmful effects of drugs and other toxicants (Ramirez et al., 2018). Therapeutic agents used against hepatic disorders possess limited therapeutic efficacy. Hence, it has become imperative to develop new, effective, safe drugs to manage liver disorders (Jin et al., 2021). The use of abundant medicinal plants and formulas with hepatoprotective efficacy to manage hepatic disorders has gained considerable attention (Ilyas et al., 2016; Jadeja et al., 2015). For example, the flavonoids extract of *Silybum marianum* plant, currently standardised as silymarin has been reported to be an effective hepatoprotective agent ((Mukhtar et al., 2021; Papackova et al., 2018; Vargas-mendoza et al., 2014). This informed the rationale for the current experiment to evaluate the effectiveness of the methanol root bark of *Acacia sieberiana* in the PCM and BDL-induced rat models of liver injury with altered biochemical parameters, oxidative stress biomarkers and liver histopathological alterations to validate its folkloric use in liver diseases.

The acute toxicity evaluation is important in checking the potentially noxious effects of chemical agents after acute administration. Besides, it helps in the LD_50_ determination of new bioactive compounds for biological screening (Kpemissi et al., 2020; Musila et al., 2017). The lack of toxic indications and death in animals following the acute administration of the compound at a dose of 5,000 mg/kg after a 14-days observation period showcases that the LD_50_ of the said compound could be higher than the 5 g/kg (Qin et al., 2009). Thus, the oral LD_50_ of the ASE in the current experiment might be greater than 5,000 mg/kg, and it could be safe at that dose via oral administration. However, mortality was observed following the *i*.*p* administration of the ASE 1,600 mg/kg resulting in a calculated *i*.*p* LD_50_ of 1,300mg/kg, as an indication that it could be relatively safe at the dose after acute *i*.*p*. administration.

Paracetamol is among the hepatotoxic therapeutic agents that cause acute hepatic failure at a high dose (Sinaga et al., 2021). Approximately one-fourth of the administered PCM binds to plasma protein and is partially biotransformed by the hepatic microsomal enzymes. It can also be metabolized by conjugation with glucuronic acid in the liver (Sinaga et al., 2021). The cytochrome P450 changes about 5-15% of PCM within the therapeutic range to a highly reactive product N-acetyl-p-benzoquinonymin (NAPQI), which is easily nullified by glutathione conjugation (Ohba et al., 2016). Following the PCM intoxication, the NAPQI targets the mitochondrial protein and interferes with the energy production, generating ROS, which leads to oxidative stresses and subsequently generates hepatotoxicity (Tsai et al., 2018; Yoon et al., 2016).

The AST, ALT, and ALP are parameters that reveal the hepatic metabolic activities (El Kabbaoui et al., 2017). The AST and ALT have been particularly elevated in conditions that cause hepatocellular damage that eventually causes leakage of these enzymes into the blood in liver injuries (Ahmad et al., 2022; Y. Li et al., 2019). The ALP level increases in case of bile duct obstruction and indicates liver or bone disorder (Sinaga et al., 2021). The effective decline in the ALT and AST levels by the ASE after the PCM hepatotoxic injury in the current work at the 250 and 750 mg/kg may be associated with its hepatoprotective efficacy. Sometimes, the toxic injury had been reported not to affect the ALP concentration, and this seemed to be confirmed in this study in that the PCM toxic injury did not significantly increase the ALP activity, unlike the ALT and AST.

The reduction in the total protein and albumin plasma concentration and the elevated levels of total and direct bilirubin in the hepatotoxic rats showcases interference of the hepatic cellular integrity and function as a result of paracetamol intoxication (Yoon et al., 2016). The ASE at all doses in this research increased the concentrations of the total protein and albumin efficiently, suggestive of its hepatoprotective activity. Besides, the ASE reduced the PCM-induced elevated bilirubin level, which was significant only in the 750 mg/kg ASE-treated group for the direct bilirubin.

The pathogenesis of hepatotoxicity is mostly related to the metabolic changes of xenobiotics to ROS, which eventually result in oxidative stress and damage to the hepatic cellular contents (Zhang et al., 2018). The antioxidant enzymes (SOD, CAT, and GPx) produce defensive mechanisms against ROS. Therefore, the imbalance between the generated ROS and antioxidant enzyme activity is responsible for the mechanisms of hepatic diseases, including hepatitis, cirrhosis, necrotic hepatitis and hepatocellular carcinoma (Banerjee et al., 2020). The SOD enzyme converts superoxide radicals to an oxygen molecule and hydrogen peroxide to maintain a steady oxygen state (Palanivel et al., 2008). CAT and GPx often convert the hydrogen peroxide into water to complete the SOD’s scavenging activity (Li et al., 2015). In this process, the toxic species (superoxide radical and hydrogen peroxide) are modified to the harmless water. Therefore, the reduction in the SOD level as seen in the PCM hepatotoxic group in this study suggests hepatocellular damage, which was efficiently attenuated in all the ASE-treated groups as an indication of protection against the PCM toxic injury. Although the PCM toxic injury did not affect the CAT concentration, ASE at 250 and 750 mg/kg as with silymarin elevated its level to indicate its hepatic boosting ability. The ASE ameliorated the injury from the PCM-induced reduction in GPx at all doses, almost to the same extent as the standard agent. It was also obvious that the PCM-induced hepatic damage results in lipid peroxidation, as evident with the enhanced MDA level in the current research. The MDA generated as a result of lipid peroxidation tend to amplify cellular damage, thus, are often used as biomarkers of oxidative liver damage (Wang et al., 2019). Hence, oxidative hepatic damage could be alleviated by inhibiting the lipid peroxidation and MDA production (Liu et al., 2015). The reduction in the MDA concentration in both the silymarin and ASE administered groups in the current work further explained the ameliorative effects of the extract against the hepatic lipid peroxidation, free radicals and cellular damage.

The reduction in the PCM-induced intense necrosis and congestion in the silymarin and ASE treated groups further indicated the hepatoprotective activity resulting probably by increasing the activity of the antioxidant system in the body, thereby preventing liver damage. This protection seemed better with the lower ASE doses than 1,500 mg/kg.

The BDL is an animal model used to induce extrahepatic cholestasis in rats accompanied by fibrosis and oxidative stress (Pereira et al., 2009). Cholestasis can result from abnormal bile secretion by the hepatic and bile duct cells or by the blockade of bile ducts (Moslemi et al., 2021). The increased concentrations of ALT, ALP, AST, total protein and bilirubin serve as sensitive markers of cholestasis to indicate bile acid degradation (Qiao et al., 2019). Generally, AST, ALT and ALP elevations are associated with hepatic damage, necrosis, and bile salts accumulation, respectively (Aryal et al., 2019; Moslemi et al., 2021). The BDL-elicited liver damage in this study seemed to suggest milder toxicity than in the PCM-induced model. Similar to the PCM-induced model, the ASE in the current experiment demonstrated ameliorative effects in the BDL-induced hepatotoxicity. All the extract groups reduced the elevated ALT concentration. However, the reduced ALT level at the lowest extract dose, which showed more of the reduction, was not significant to the BDL-injury. The BDL-injury elevated AST was also reduced in all the extract groups, but only the lower extract doses were significantly reduced. However, ALP which was also reduced in all the treatment groups, was only significant at the higher extract doses. The better result exhibited by the lower doses of the extract for ALT and AST seemed to suggest its liver protection.

The effect of ligation was slight for both total protein (which increased) and albumin (decreased slightly). The ASE at the higher doses increased the albumin level, but total protein slightly decreased in all the ASE doses, which were insignificant. The direct and total bilirubin elevated with the BDL injury were significantly reduced in all the extract-treated groups, which could be related to the enhanced bilirubin uptake or its conjugation by the extract (Zhu et al., 2018). Therefore, the ASE in the present work might have increased the bile formation as part of its hepatoprotective efficacy.

Oxidative stress is the major pathogenesis in cholestasis, resulting from an imbalance between the antioxidant and oxidative systems (Moslemi et al., 2021). Some enzymes, such as CAT, SOD, and GPx, are antioxidants (Moslemi et al., 2021). The decrease in SOD from BDL-injury was prevented in the lower ASE doses (125 and 250 mg/kg) better than the silymarin. Although the SOD level was increased in all the treatment groups, the level was still significantly below that of the normal control. A dose-dependent increase in the CAT concentration also occurred with all the ASE-treated groups, and in this case, the increase at the 380 mg/kg extract dose was more and was not significantly different from the silymarin group unlike the lower extract doses. In both the PCM- and BDL-induced injury, the GPx concentration was not significantly reduced, but it was increased in all the treatment groups of both the standard drug and ASE dose groups. GPx is an enzyme that protects haemoglobin from oxidative degradation in the red blood cells (Yoshikawa & Naito, 2002). The enzyme has been reported to be over-expressed to protect cells against oxidative damage and apoptosis induced by hydrogen peroxide (Yoshikawa & Naito, 2002). It was also evident that the lowest ASE dose reduced the BDL-induced elevated MDA level, suggesting protection against the oxidative free radical, hepatic lipid peroxidation, and hepatocellular injury.

The moderate necrosis and vascular congestion observed with the BDL-produced liver damage were reversed by the 125mg/kg extract group. Besides, milder effects were observed in the higher ASE dose groups (250 and 380 mg/kg). These effects could suggest some level of liver protection and regenerative potential of the extract.

The therapeutic potentials of the medicinal plant are usually a direct function of the bioactive components in the plant, which could serve as a lead to develop hepatoprotective compounds (Girish et al., 2009). The present experiment showcased that glycosides, triterpenes, saponins, tannins, flavonoids and alkaloids could be available in the ASE. These compounds mostly act as antioxidants to abolish free radical generation and eventually prevent hepatic damage (Ramadan et al., 2013). For instance, flavonoids compounds such as catechin and kaempferol fight against free radicals and inhibit liver disease pathogenesis (Okaiyeto et al., 2018). Also, glycosides act as an antioxidant to provide hepatoprotective efficacy (Asaad et al., 2021; Siddiqui et al., 2018). Besides, (Huang et al., 2014) reported the hepatic protection of triterpenoids and saponins. A previous study by (Jan et al., 2017) has shown that alkaloids reduced biochemical biomarkers (ALT, AST, and ALP), attenuated hepatic inflammation, and elevated the antioxidant enzyme concentration in liver damage. Hence, the hepatic protective potentials of the ASE observed in the current research could be related to the presence of these phytocomponents.

## 5.0 Conclusion

The research findings have shown that the root bark extract of *Acacia sieberiana* ameliorates chemically-induced and cholestatic liver damages, possibly via modulating the biochemical and oxidative stress biomarkers, which justifies its ethnomedicinal value against hepatic disorders.

## List of abbreviations

ABU: Ahmadu Bello University
ABUCAUC: ABU Ethical Committee on Animal Use and Care Research Policy
ALP: Alkaline phosphatase
ALT: Alanine transaminase
NOVA: Analysis of variance
ARRIVE: Animal Research: Reporting of *In Vivo* Experiments
ASE: *Acacia sieberiana* extract
AST: aspartate transaminase
BDL: Bile duct ligation
CAT: Catalase
DB: Direct bilirubin
DW: Distilled water
FN: Focal necrosis
GPx: Glutathione peroxidase
*i*.*p*: Intraperitoneal
KH: Kupffer cell hyperplasia
LD50: Medial lethal dose
LH: Lymphocytes hyperplasia
MDA: Malondialdehyde
MN: Moderate necrosis
N: Necrosis
NAPQI: N-acetyl-p-benzoquinonymin
NH: Normal hepatocytes
PCM: Paracetamol
ROS: Reactive oxygen species
rpm: Revolutions per minute
SC: Sinusoid congestion
SC: slight sinusoidal congestion
SEM: Standard error of the mean
SN: Slight necrosis
SOD: Superoxide dismutase
TB: Total bilirubin
TP: Total protein
VC: Vascular congestion

## Declaration

### Ethics approval and consent to participate

The protocols for the research were approved by the Ahmadu Bello University Ethical Committee on Animal Use and Care Research Policy (permission number: ABUCAUC/2016/049) and conducted based on the Animal Research: Reporting of *In Vivo* Experiments (ARRIVE) guidance.

### Consent for publication

Not applicable

### Availability of data and material

The datasets generated during and/or analyzed during the experimental work are available from the corresponding author on reasonable request

### Authors’ Contribution

**MW:** Conceptualization, resources, investigation, writing original draft, data curation, review, editing, and data analysis. **JIE:** Supervision, validation, review, and project administration. **AM:** Investigation, review, and editing. **MHA**: Investigation, writing the original draft, revised and edit the entire manuscript. All the authors read and approved the final version of the manuscript

## Acknowledgements

The authors appreciate the entire staff of the Department of Pharmacology and Therapeutics, ABU, Zaria, Nigeria, for their support throughout the period of the experiment.

## References

Abdelhady, M. I. (2013). A novel polyphenolic compound isolated from Acacia sieberiana. Organic Chemistry, 9(6), 236–238.

Abenavoli, L., Izzo, A. A., Natasa, M., Cicala, C., Santini, A., & Capasso, R. (2018). Milk thistle (Silybum marianum): A concise overview on its chemistry, pharmacological, and nutraceutical uses in liver diseases. Phytotherapy Research, 1–12. https://doi.org/10.1002/ptr.6171

Ahmad, M. H., Zezi, A. U., Bola, A. S., Alhassan, Z., Mohammed, M., & Danraka, R. N. (2020). Mechanisms of antidiarrhoeal activity of methanol leaf extract of Combretum hypopilinum Diels (Combretaceae): Involvement of opioidergic and (α1 and β) –adrenergic pathways. Journal of Ethnopharmacology, 269, 113750. https://doi.org/10.1016/j.jep.2020.113750

Ahmad, M. H., Zezi, A. U., Sherifat, B. A., Alshargi, O. Y., Mohammed, M., Mustapha, S., Bala, A. A., Muhammad, S., Julde, S. M., Wada, A. S., & Jatau, I. A. (2022). Sub-acute toxicity study on hydromethanolic leaves extract of Combretum hypopilinum (Combretaceae) Diels in Wistar rats. Toxicological Research, 0123456789, 1–16. https://doi.org/10.1007/s43188-022-00133-5

Ali, S. A., Sharief, N. H., & Mohamed, Y. S. (2019). Hepatoprotective Activity of Some Medicinal Plants in Sudan. Evidence-Based Complementary and Alternative Medicin, 2196315, 1–16. https://doi.org/10.1155/2019/2196315

Review Ameh, P. O., & Eddy, N. O. (2014). Characterisation of Acacia Sieberiana (AS) Gum and Their Corrosion Inhibition Potentials for Zinc in Sulphuric Acid Medium. International Journal of Novel Research in Physics Chemistry & Mathematics, 1(1), 25–36.

Anyasor, G. N., Moses, N., & Kale, O. (2020). Hepatoprotective and hematological effects of Justicia secunda Vahl leaves on carbon tetrachloride-induced toxicity in rats. Biotechnic & Histochemistry, 00(00), 1–11. https://doi.org/10.1080/10520295.2019.1700430

Aryal, S., Adhikari, B., Panthi, K., Aryal, P., Sharifi-rad, J., & Koirala, N. (2019). Antipyretic, Antinociceptive, and Anti-InflammatorActivities from Pogostemon benghalensis Leaf Extract in Experimental Wister Rats. Medicines, 6, 96. https://doi.org/doi:10.3390/medicines6040096

Asaad, G. F., Mohammed, H., Abdallah, I., Mohammed, H. S., & Nomier, Y. A. (2021). Hepatoprotective effect of kaempferol glycosides isolated from Cedrela odorata L. leaves in albino mice: involvement of Raf/MAPK pathway. Research in Pharmaceutical Sciences, 16(4), 370–380.

Asrani, S. K., Devarbhavi, H., Eaton, J., & Kamath, P. S. (2019). Burden of liver diseases in the world. Journal of Hepatology, 70(1), 151–171. https://doi.org/10.1016/j.jhep.2018.09.014

Banerjee, A., Das, D., Paul, R., Roy, S., Bhattacharjee, A., Ug, P., College, S., Road, W. C., & Bengal, W. (2020). Altered composition of high-lipid diet may generate reactive oxygen species by disturbing the balance of antioxidant and free radicals: Journal of Basic and Clinical Physiology and Pharmacology, 20190141, 1–19. https://doi.org/10.1515/jbcpp-2019-0141

Bernal, W., Auzinger, G., Dhawan, A., & Wendon, J. (2010). Acute liver failure. The Lancet, 376(9736), 190–201. https://doi.org/10.1016/S0140-6736(10)60274-7

Dawurung, C. J., Elisha, I. L., Offiah, N. V, Gotep, J. G., Oladipo, O. O., Makoshi, M. S., Makama, S., & Shamaki, D. (2012). Antidiarrheal Evaluation of Aqueous and Ethanolic Leaf Extracts of Acacia sieberiana DC. (Fabaceae) in Albino Rats. Asian Journal of Experimental Biological Sciences, 4(4), 799–803.

Duncan, A. W., Dorrell, C., & Grompe, M. (2009). Stem Cells and Liver Regeneration. Gastroenterology, 137(2), 466–481. https://doi.org/10.1053/j.gastro.2009.05.044

El Kabbaoui, M., Chda, A., El-Akhal, J., Azdad, O., Mejrhit, N., Aarab, L., Bencheikh, R., & Tazi, A. (2017). Acute and sub-chronic toxicity studies of the aqueous extract from leaves of Cistus ladaniferus L. in mice and rats. Journal of Ethnopharmacology, 209, 147–156. https://doi.org/10.1016/j.jep.2017.07.029

Elmasry, S. (2021). The hepatoprotective effect of gooseberry and black mulberry extracts against carbon tetrachloride-induced liver injury in rats. The Journal of Basic and Applied Zoology, 82(33), 1–11. https://doi.org/10.1186/s41936-021-00224-z

Elufioye, T. O., & Habtemariam, S. (2019). Hepatoprotective e ff ects of rosmarinic acid: Insight into its mechanisms of action. Biomedicine & Pharmacotherapy Journal, 112, 1–11. https://doi.org/10.1016/j.biopha.2019.108600

Girish, C., Koner, B. C., Jayanthi, S., Rao, K. R., Rajesh, B., & Pradhan, S. C. (2009). Hepatoprotective activity of picroliv, curcumin and ellagic acid compared to silymarin on paracetamol induced liver toxicity in mice. Fundamental & Clinical Pharmacology, 23, 735–745. https://doi.org/10.1111/j.1472-8206.2009.00722.x

Huang, B., Fu, H., Chen, W., Luo, Y., & Ma, S. (2014). Hepatoprotective Triterpenoid Saponins from Callicarpa nudiflora. Chemical and Pharmaceutical Bulletin, 62(July), 695–699. https://doi.org/10.1248/cpb.c14-00159

Ilyas, U., Katare, D. P., Aeri, V., & Naseef, P. P. (2016). Review on Hepatoprotective and Immunomodulatory Herbal Plants. Pharmacognosy Review, 10(19), 66–70. https://doi.org/10.4103/0973-7847.176544

Jadeja, R., Devkar, R. V, & Nammi, S. (2015). Hepatoprotective Potential of Herbal Medicine. Evidence-Based Complementary and Alternative Medicine, 536564. http://dx.doi.org/10.1155/2015/536564

Jan, N. U., Ahmad, B., Ali, S., Adhikari, A., Ali, A., Jahan, A., Ali, A., & Ali, H. (2017). Steroidal Alkaloids as an Emerging Therapeutic Alternative for Investigation of Their Immunosuppressive and Hepatoprotective Potential. Frontiers in Pharmacology, 8, 1–13. https://doi.org/10.3389/fphar.2017.00114

Jin, S. E., Shin, H., & Ha, H. (2021). Hepatoprotective effects of Gamisoyo-san against acetaminophen-induced liver injuries. Integrative Medicine Research, 10(1), 100466. https://doi.org/10.1016/j.imr.2020.100466

Khan, A., Najeeb-ur-Rehman Alkharfy, K. M., & Gilani, A. H. (2011). Antidiarrheal and antispasmodic activities of Salvia officinalis are mediated through activation of K + channels. Bangladesh Journal of Pharmacology, 6(2), 111–116. https://doi.org/10.3329/bjp.v6i2.9156

Khan, H., Ullah, H., & Nabavi, S. M. (2019). Mechanistic insights of hepatoprotective effects of curcumin: Therapeutic updates and future prospects. Food and Chemical Toxicology. https://doi.org/10.1016/j.fct.2018.12.002

Kpemissi, M., Metowogo, K., Melila, M., Veerapur, V. P., Negru, M., Taulescu, M., Potârniche, A., Shivalingaiah, D., Adinarayanashetty, T., Vijayakumar, S., Eklu-gadegbeku, K., & Aklikokou, K. (2020). Acute and subchronic oral toxicity assessments of Combretum micranthum (Combretaceae) in Wistar rats. Toxicology Reports, 7, 162–168. https://doi.org/10.1016/j.toxrep.2020.01.007

Li, S., Tan, H., Wang, N., Zhang, Z., Lao, L., Wong, C., & Feng, Y. (2015). The Role of Oxidative Stress and Antioxidants in Liver Diseases. International Journal of Molecular Sciences, 16(11), 26087–26124. https://doi.org/10.3390/ijms161125942

Li, Y., Kandhare, A. D., Mukherjee, A. A., & Bodhankar, S. L. (2019). Acute and sub-chronic oral toxicity studies of hesperidin isolated from orange peel extract in Sprague Dawley rats. Regulatory Toxicology and Pharmacology, 105(1), 77–85. https://doi.org/10.1016/j.yrtph.2019.04.001

Lin, S., Wang, Y., Chen, W., Liao, S., Chou, S., Yang, C., & Chen, C. (2017). Hepatoprotective activities of rosmarinic acid against extrahepatic cholestasis in rats. Food and Chemical Toxicology, 108, 214–223. https://doi.org/10.1016/j.fct.2017.08.005

Liu, J., Lu, J., Wen, X., Kan, J., & Jin, C. (2015). Antioxidant and protective effect of inulin and catechin grafted inulin against CCl 4 -induced liver injury. International Journal of Biological Macromolecules, 72, 1479–1484. https://doi.org/10.1016/j.ijbiomac.2014.09.066

Lorke, D. (1983). A New Approach to Practical Acute Toxicity Testing. Archives of Toxicology, 54, 275–287.

Ma, X., Jiang, Y., Zhang, W., Wang, J., Wang, R., Wang, L., Wei, S., Wen, J., Li, H., & Zhao, Y. (2020). Natural products for the prevention and treatment of cholestasis: A review. Phytotherapy Research, 34(6), 1291–1309. https://doi.org/10.1002/ptr.6621

Madrigal-santillán, E., Madrigal-bujaidar, E., Álvarez-gonzález, I., Sumaya-martínez, M. T., Gutiérrez-salinas, J., Bautista, M., Morales-gonzález, Á., González-rubio, M. G., Aguilar-faisal, J. L., Morales-gonzález, J. A., Madrigal-bujaidar, E., & Álvarez-gonzález, I. (2014). Review of natural products with hepatoprotective effects. World Journal of Gastroenterology, 20(40), 14787–14804. https://doi.org/10.3748/wjg.v20.i40.14787

Mahmood, N. D., Mamat, S. S., Kamisan, F. H., Yahya, F., Kamarolzaman, M. F. F., Nasir, N., Mohtarrudin, N., Tohid, S. F., & Zakaria, Z. A. (2014). Amelioration of Paracetamol-Induced Hepatotoxicity in Rat by the Administration of Methanol Extract of Muntingia calabura L. Leaves. BioMed Research International, 2014(695678). http://dx.doi.org/10.1155/2014/695678

Michalopoulos, G. K. (2017). Hepatostat: Liver Regeneration and Normal Liver Tissue Maintenance. Hepatology, 65(4), 1384–1392. https://doi.org/10.1002/hep.28988

Miltonprabu, S., Tomczyk, M., Skalicka-Woźniak, K., Rastrelli, L., Daglia, M., Nabavi, S. F., Alavian, S. M., & Seyed, N. S. M. (2016). Hepatoprotective effect of quercetin: From chemistry to medicine. Food and Chemical Toxicology. https://doi.org/10.1016/j.fct.2016.08.034

Moslemi, Z., Bahrami, M., Hosseini, E., Mansourian, M., Daneshyar, Z., Eftekhari, M., Shakerinasab, N., Asfaram, A., Kokhdan, E. P., Barmoudeh, Z., & Doustimotlagh, A. H. (2021). Portulaca oleracea methanolic extract attenuate bile duct ligation-induced acute liver injury through hepatoprotective and anti-inflammatory effects. Heliyon, 7, e07604. https://doi.org/10.1016/j.heliyon.2021.e07604

Muhammad, S. L., Wada, Y., Mohammed, M., Ibrahim, S., Musa, K. Y., Olonitola, O. S., Ahmad, M. H., Mustapha, S., Rahman, Z. A., & Sha’aban, A. (2021). Bioassay-Guided Identification of Bioactive Compounds from Senna alata L. against Methicillin-Resistant Staphylococcus aureus. Applied Microbiology, 1, 520–536. https://doi.org/10.3390/applmicrobiol1030034

Mukhtar, S., Xiaoxiong, Z., Qamer, S., Saad, M., Mubarik, M. S., Mohammed, A. H., & Mohammed, O. B. (2021). Hepatoprotective activity of silymarin encapsulation against hepatic damage in albino rats. Saudi Journal of Biological Sciences, 28(1), 717–723. https://doi.org/10.1016/j.sjbs.2020.10.063

Musila, M. N., Ngai, D. N., Mbiri, J. W., Njagi, S. M., Mbinda, W. M., & Ngugi, M. P. (2017). Acute and Sub-Chronic Oral Toxicity Study of Methanolic Extract of Caesalpinia volkensii (Harms). Journal of Drug Metabolism & Toxicology, 08(01), 1–8. https://doi.org/10.4172/2157-7609.1000222

Ngaffo, C. M. N., Tchangna, R. S. V, Mbaveng, A. T., Kamga, J., Harvey, F. M., Ngadjui, B. T., Bochet, C. G., & Kuete, V. (2020). Botanicals from the leaves of Acacia sieberiana had better cytotoxic effects than isolated phytochemicals towards MDR cancer cells lines. Heliyon, 6, e05412. https://doi.org/10.1016/j.heliyon.2020.e05412

Ohba, H., Kanazawa, M., Kakiuchi, T., & Tsukada, H. (2016). Effects of acetaminophen on mitochondrial complex I activity in the rat liver and kidney: a PET study with 18 F-BCPP-BF. EJNMMI Research, 682, 1–10. https://doi.org/10.1186/s13550-016-0241-4

Ohemu, T. L., Agunu, A., Olotu, P. N., Ajima, U., Dafam, D. G., & Azila, J. J. (2014). Ethnobotanical survey of medicinal plants used in the traditional treatment of viral infections in Jos, Plateau state, Nigeria. International Journal of Medicinal and Aromatic Plants, 4(2), 74–81.

Okaiyeto, K., Nwodo, U. U., Mabinya, L. V, & Okoh, A. I. (2018). A Review on Some Medicinal Plants with Hepatoprotective Effects. Pharmacognosy Review, 12(24), 186–199. https://doi.org/10.4103/phrev.phrev

Palanivel, M. G., Rajkapoor, B., Kumar, R. S., Einstein, J. W., Kumar, E. P., Kumar, M. R., Kunchu, K., Pradeep, K. M., & Balasundaram, J. (2008). Hepatoprotective and Antioxidant Effect of Pisonia aculeata L. against CCl4-Induced Hepatic Damage in Rats. Scientia Pharmaceutica, 76(2), 203–215. https://doi.org/10.3797/scipharm.0803-16

Papackova, Z., Heczkova, M., Dankova, H., Sticova, E., Lodererova, A., Bartonova, L., Poruba, M., & Cahova, M. (2018). Silymarin prevents acetaminophen-induced hepatotoxicity in mice. PLoS ONE, 13(1), e019135. https://doi.org/10.1371/journal.pone.0191353

Park, H., Jo, E., Han, J., Jung, S., Lee, D., Park, I., Heo, K., Na, M., & Myung, C. (2019). Hepatoprotective effects of an Acer tegmentosum Maxim extract through antioxidant activity and the regulation of autophagy. Journal of Ethnopharmacology, 239, 111912. https://doi.org/10.1016/j.jep.2019.111912

Pereira, F. A., Facincan, I., Jorgetti, V., Ramalho, L. N. Z., Volpon, J. B., Reis Dos, L. M., & Paula, F. J. A. (2009). Etiopathogenesis of Hepatic Osteodystrophy in Wistar Rats with Cholestatic Liver Disease. Calcified Tissue International, 85(1), 75–83. https://doi.org/10.1007/s00223-009-9249-3

Pingili, R. B., Pawar, A. K., Challa, S. R., Kodali, T., Koppula, S., & Toleti, V. (2019). A comprehensive review on hepatoprotective and nephroprotective activities of chrysin against various drugs and toxic agents. Chemico-Biological Interactions, 308, 51–60. https://doi.org/10.1016/j.cbi.2019.05.010

Protzer, U., Maini, M. K., & Knolle, P. A. (2012). Living in the liver: hepatic infections. Nature Reviews Immunology, 12(3), 201–213. https://doi.org/10.1038/nri3169

Qiao, J., Li, H., Liu, F., Li, Y., Tian, S., & Cao, L. (2019). Effects of Portulaca oleracea Extract on Acute Alcoholic Liver Injury of Rats. Molecules, 24(16), 2887. https://doi.org/10.3390/molecules24162887

Qin, Y., Wu, X., Huang, W., Gong, G., Li, D., He, Y., & Zhao, Y. (2009). Acute toxicity and sub-chronic toxicity of steroidal saponins from Dioscorea zingiberensis C. H. Wright in rodents. Journal of Ethnopharmacology Journal, 126, 543–550. https://doi.org/10.1016/j.jep.2009.08.047

Ramadan, M. M., Abd-algader, N. N., El-kamali, H. H., Ghanem, K. Z., & Farrag, A. R. (2013). Volatile compounds and antioxidant activity of the aromatic herb Anethum graveolens. Journal of the Arab Society for Medical Research, 8(2), 79–88. https://doi.org/10.4103/1687-4293.123791

Ramirez, T., Strigun, A., Verlohner, A., Huener, H.-A., Peter, E., Herold, M., Bordag, N., Mellert, W., Walk, T., Spitzer, M., Jiang, X., Sperber, S., Hofmann, T., Hartung, T., Kamp, H., & Ravenzwaay, B. Van. (2018). Prediction of liver toxicity and mode of action using metabolomics in vitro in HepG2 cells. Archives of Toxicology, 92(2), 893–906. https://doi.org/10.1007/s00204-017-2079-6

Rašković, A., Milanovi, I., Pavlović, N., Ćebović, T., Vukmirović, S., & Mikov, M. (2014). Antioxidant activity of rosemary (Rosmarinus officinalis L.) essential oil and its hepatoprotective potential. BMC Complementary and Alternative Medicine, 14(225), 1–9. https://doi.org/10.1186/1472-6882-14-225

Salisu, A., Ogbadu, G. H., Onyenekwe, P. C., Olorode, O., Ndana, R. W., & Segun, O. (2014). Evaluating the Nutritional Potential of Acacia Sieberiana Seeds (Dc) Growing in North West of Nigeria. Journal of Biology and Life Sciences, 5(2), 26–36.

Seigler, D. S., Butterfield, C. S., Dunn, J. E., & Conn, E. E. (1975). Dihydroacacipetalin–a new cyanogenic glucoside from Acacia sieberiana var. Woodi. Phytochemistry, 14, 1419–1420.

Siddiqui, M. A., Ali, Z., Chittiboyina, A. G., & Khan, I. A. (2018). Hepatoprotective Effect of Steroidal Glycosides From Dioscorea villosa on Hydrogen Peroxide-Induced Hepatotoxicity in HepG2 Cells. Frontiers in Pharmacology, 9, 1–12. https://doi.org/10.3389/fphar.2018.00797

Sinaga, E., Fitrayadi, A., Asrori, A., Rahayu, S. E., Suprihatin, S., & Prasasty, V. D. (2021). Hepatoprotective effect of Pandanus odoratissimus seed extracts on paracetamol-induced rats induced rats. Pharmaceutical Biology, 59(1), 31–39. https://doi.org/10.1080/13880209.2020.1865408

Sofowora, A. (1993). Medicinal Plants and Traditional Medicine in Africa (2nd Editio). Spectrum Books Ltd.

Tag, C. G., Sauer-lehnen, S., Weiskirchen, S., Borkham-kamphorst, E., Tolba, R. H., Tacke, F., & Weiskirchen, R. (2015). Bile Duct Ligation in Mice: Induction of Inflammatory Liver Injury and Fibrosis by Obstructive Cholestasis. Journal of Visualized Experiments, 96(e52438), 1–11. https://doi.org/10.3791/52438

Tsai, M.-S., Wang, Y.-H., Lai, Y.-Y., Tsou, H., Liou, G., Ko, J., & Wang, S.-H. (2018). Kaempferol protects against propacetamol-induced acute liver injury through CYP2E1 inactivation, UGT1A1 activation, and attenuation of oxidative stress, inflammation and apoptosis in mice. Toxicology Letters, 290, 97–109. https://doi.org/10.1016/j.toxlet.2018.03.024

Usman, A. M., Danjuma, N. M., Ya’u, J., Ahmad, M. M., Alhassan, Z., Abubakar, Y. M., & Ahmad, M. H. (2021). Antidiarrhoeal potentials of methanol bark extract of Hymenocardia Acida Tul (Euphorbiaceae) in laboratory animals. Bulletin of the National Research Centre, 45(118), 1–12. https://doi.org/10.1186/s42269-021-00575-1

Uzunhisarcikli, M., & Aslanturk, A. (2019). Hepatoprotective effects of curcumin and taurine against bisphenol A-induced liver injury in rats. Environmental Science and Pollution Research, 26(36), 37242–37253. https://doi.org/10.1007/s11356-019-06615-8

Vargas-mendoza, N., Madrigal-santillán, E., Morales-gonzález, Á., Esquivel-soto, J., Esquivel-chirino, C., González-rubio, M. G., Gayosso-de-lucio, J. A., & Morales-gonzález, J. A. (2014). Hepatoprotective effect of silymarin. World Journal of Hepatology, 6(3), 144–149. https://doi.org/10.4254/wjh.v6.i3.144

Wang, Y. A. O., Chen, Q., Shi, C., Jiao, F., & Gong, Z. (2019). Mechanism of glycyrrhizin on ferroptosis during acute liver failure by inhibiting oxidative stress. Molecular Medicine Reports, 20, 4081–4090. https://doi.org/10.3892/mmr.2019.10660

Yoon, E., Babar, A., Choudhary, M., Kutner, M., & Pyrsopoulos, N. (2016). Acetaminophen-Induced Hepatotoxicity: a Comprehensive Update. Journal of Clinical and Translational Hepatology, 4(2), 131–142. https://doi.org/10.14218/JCTH.2015.00052

Yoshikawa, T., & Naito, Y. (2002). What Is Oxidative Stress_□? Japan Medical Association Journal, 124(11), 271–276.

Younossi, Z. M., Stepanova, M., Afendy, M., Fang, Y., Younossi, Y., Mir, H., & Srishord, M. (2011). Changes in the Prevalence of the Most Common Causes of Chronic Liver Diseases in the United States From 1988 to 2008. Clinical Gastroenterology and Hepatology, 9(6), 524–530. https://doi.org/10.1016/j.cgh.2011.03.020

Yue, S., Xue, N., Li, H., Huang, B., Chen, Z., & Wang, X. (2020). Hepatoprotective Effect of Apigenin Against Liver Injury via the Non-canonical NF-κ B Pathway In Vivo and In Vitro. Inflammation, 43(5), 1634–1648. https://doi.org/10.1007/s10753-020-01238-5

Zhang, C., Wang, N., Xu, Y., Tan, H., Li, S., & Feng, Y. (2018). Molecular Mechanisms Involved in Oxidative Stress-Associated Liver Injury Induced by Chinese Herbal Medicine_□: An Experimental Evidence-Based Literature Review and Network Pharmacology Study. International Journal of Molecular Sciences, 19(9), 2745. https://doi.org/10.3390/ijms19092745

Zhou, H., Wang, H., Shi, K., Li, J., Zong, Y., & Du, R. (2019). Hepatoprotective Effect of Baicalein Against Acetaminophen-Induced Acute Liver Injury in Mice. Molecules, 24(1), 131. https://doi.org/10.3390/molecules24010131

Zhu, L., Wang, L., Cao, F., Liu, P., Bao, H., Yan, Y., Dong, X., Dong, W., Wang, Z., & Gong, P. (2018). Modulation of transport and metabolism of bile acids and bilirubin by chlorogenic acid against hepatotoxity and cholestasis in bile duct ligation rats: involvement of SIRT1-mediated deacetylation of FXR and PGC-1. Journal of Hepato-Biliary-Pancreatic Sciences, 25(3), 195–205. https://doi.org/10.1111/ijlh.12426

